# Household LED Light-Mediated Photodynamic Therapy for Effective Inhibition of Multidrug-Resistant *Staphylococcus aureus*

**DOI:** 10.1101/2025.08.09.668366

**Authors:** Lama Misba, Farheen Akhtar, Sameera Mujahid, Asad U. Khan

## Abstract

This study investigates the efficacy of toluidine blue O (TBO) and curcumin as photosensitizers for photodynamic therapy (PDT) against *Staphylococcus aureus* using a household LED bulb as the light source. Antibacterial and anti-biofilm activities were evaluated via colony-forming unit (CFU) and Congo red assays, while reactive oxygen species (ROS) generation was quantified using DCFH-DA dye. Confocal laser scanning microscopy (CLSM) was employed to assess live/dead bacterial cell ratios, and an *in vivo* skin abrasion model in Wistar rats was used to validate therapeutic outcomes. The CFU assay revealed substantial bacterial reductions, achieving 4.85 log₁₀ CFU/ml at 10 minutes for TBO and 2.81 log₁₀ CFU/ml at 15 minutes for curcumin. Extracellular polymeric substance (EPS) production decreased by 96.22% with TBO and 48.98% with curcumin, accompanied by enhanced ROS generation. CLSM confirmed a higher dead-to-live cell ratio following PDT. *In vivo* results, supported by histopathological examination and cytokine expression profiling, further demonstrated the effectiveness of TBO- and curcumin-mediated PDT under LED light. Overall, these findings highlight the potential of readily available household LED bulbs as an accessible and efficient light source for PDT, offering robust antimicrobial activity against *S. aureus*.

## INTRODUCTION

Antimicrobial resistance poses a significant and often overlooked threat, with the potential to escalate into a global health crisis if not addressed promptly (Murray et al. 2022; Ikuta et al. 2022). A report published by the Centers for Disease Control and Prevention (CDC) in July 2024, *Antimicrobial Resistance Threats in the United States, 2021-2022*, highlighted a 20% increase in hospital-acquired infections caused by six antimicrobial-resistant bacterial species following the COVID-19 pandemic (2019). The data further revealed a concerning rise in resistant infections during hospitalization, particularly among carbapenem-resistant *Acinetobacter*, ESBL-producing *Enterobacterales*, vancomycin-resistant *Enterococcus*, multidrug-resistant *Pseudomonas aeruginosa*, and methicillin-resistant *Staphylococcus aureus* (2022). These multidrug-resistant (MDR) pathogens are responsible for common nosocomial infections, including sepsis, ventilator-associated pneumonia, lung infections, and urinary tract infections (Mazzariol et al. 2017; Minasyan et al. 2017; Singh et al. 2020). The majority of bacterial infections are attributed to MDR strains such as *Staphylococcus aureus, Pseudomonas aeruginosa, Escherichia coli, Klebsiella pneumoniae*, and *Acinetobacter baumannii* (Terreni et al. 2021; Venkateswaran et al. 2023; Denissen et al. 2022). Additionally, the ability of many bacterial strains to form biofilms further enhances their resistance to antibiotics (Sharma et al. 2019; Liu et al. 2024; Dincer et al. 2020).

The growing threat of antimicrobial resistance has underscored the need for alternative therapeutic approaches. One such strategy is photodynamic therapy (PDT), which is a safe, non-invasive, and broad-spectrum antimicrobial method that does not target specific cellular components (Mathur et al. 2023). PDT employs a photosensitizer (PS) in conjunction with a light source, where the PS, upon exposure to a specific wavelength of light in an oxygen-rich environment, undergoes excitation. This transition from the ground singlet state to a stable, long-lived excited triplet state leads to the generation of reactive oxygen species (ROS) through two distinct reaction pathways. In type 1 reactions, the excited triplet state interacts with a substrate, producing free radicals such as hydroxyl radicals (OH•) and hydrogen peroxide (H₂O₂). In type 2 reactions, the excited triplet state reacts with molecular oxygen, generating singlet oxygen, both of which are highly toxic and contribute to microbial cell death (Correia et al. 2021).

Photosensitizers are categorized into four main classes: synthetic dyes (e.g., new methylene blue, toluidine blue O), tetra-pyrrole structures (e.g., porphyrins, chlorins), natural PSs (e.g., curcumin, riboflavin), and nanostructures (e.g., fullerenes, titanium dioxide) (Ghorbani et al. 2018; Escudero et al. 2021). Synthetic dyes, possessing a cationic charge, effectively bind to both Gram-positive and Gram-negative bacteria (Abrahamse et al. 2016). Tetra-pyrrole compounds, often termed the *pigments of life*, represent some of the earliest known PSs and are capable of absorbing light at higher wavelengths (Pemula et al. 2021). Natural PSs are widely available, exhibit low cytotoxicity, and are considered safe for use, making them promising candidates for antimicrobial applications (Aebisher et al. 202). Nanostructures help overcome challenges such as hydrophobicity and aggregation, which are common issues associated with many PSs (Abrahamse et al. 2016).

In this study, we evaluated the efficacy of two PSs from distinct classes—toluidine blue O (a synthetic dye) and curcumin (a natural PS)—using a household LED bulb as the light source. Our primary objective was to assess their antimicrobial potential under LED light exposure and compare their effectiveness. Additionally, to validate our findings, we employed an *in vivo* infection model to analyse the impact of TBO and curcumin on skin abrasions in male Wistar rats following LED exposure.

## METHODS

### Ethical Statement

The *in vivo* study was conducted in compliance with the Institutional Animal Ethics Committee (IAEC) guidelines and was approved by the Institutional Animal Ethics Committee of Jawaharlal Nehru Medical College, AMU (Registration No. 401/GO/Re/S/2001/CPCSEA).

### Bacterial Strain and Culture Conditions

*Staphylococcus aureus* strain was preserved at −80 °C and routinely cultured in Brain Heart Infusion (BHI) broth (HiMedia Laboratories, Mumbai, India) at 37 °C prior to experimentation.

### Lightsource

A commonly available white light-emitting diode (LED) bulb (Philips warm and cool light variants), typically used in household and hospital settings, was employed as the light source for all experiments.

### Assessment of Planktonic Cell Inactivation by Photosensitizers Using CFU Assay

An overnight culture of *Staphylococcus aureus* was harvested, washed, and resuspended in fresh medium to achieve a final cell density of 10⁶ CFU/mL. A volume of 100 µL of this suspension was dispensed into each well of a 96-well microtiter plate and incubated at 37 °C for 24 hours to allow biofilm formation. Following incubation, non-adherent cells were removed, and the wells were gently washed twice with phosphate-buffered saline (PBS, pH 7.4) to retain only the adhered biofilm. The preformed biofilms were then incubated with 100 µM of photosensitizer (TBO or curcumin) in the dark for 30 minutes. Post-incubation, the wells were exposed to white LED light from household bulbs for 5, 10, or 15 minutes. Control groups included: (i) biofilms treated with light only (no PS), (ii) biofilms treated with PS only (no light, to assess dark toxicity), and (iii) untreated biofilms (PBS only). After treatment, the biofilms were disrupted, and the resulting suspensions were serially diluted 10-fold. A 100 µL aliquot from each dilution was plated onto Brain Heart Infusion (BHI) agar and incubated at 37 °C for 24 hours. Colony-forming units (CFUs) were then counted to assess bacterial viability (Akhtar et al. 2021).

### Evaluation of Extracellular Polymeric Substances (EPS) Production by Photosensitizers (PS)

EPS production was evaluated using the Congo Red (CR) binding assay, as previously described (Misba et al., 2018). Briefly, 100 μL of *Staphylococcus aureus* overnight culture (10⁶ CFU/mL) was added to each well of a 96-well microtiter plate and incubated at 37 °C for 24 hours to allow biofilm formation. After incubation, non-adherent cells were removed, and the wells were gently washed twice with phosphate-buffered saline (PBS, pH 7.4) to retain only the adherent biofilm. The preformed biofilms were then treated with 100 µM of each photosensitizer (TBO and curcumin) in the dark for 30 minutes, followed by exposure to white household LED light bulbs for 5, 10, or 15 minutes. Control groups included untreated biofilms and biofilms treated with PS alone (without light exposure). After treatment, the medium was removed, and the wells were gently washed with PBS. To assess EPS production, 100 μL of fresh medium and 50 μL of 0.5 mM Congo Red solution were added to each treated and control well. For blank measurements, 100 μL of medium and 50 μL of CR were added to fresh wells without cells. The plates were then incubated at 4 °C for 2 hours. Following incubation, the contents of each well were vortexed and transferred to 200 μL microcentrifuge tubes, then centrifuged at 10,000× g for 5 minutes. The resulting supernatants were transferred to a clean 96-well plate, and absorbance was measured at 490 nm using a microplate reader (Bio-Rad iMark™). EPS production was quantified using the formula:

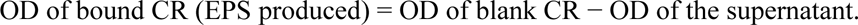

### Detection of Intracellular Reactive Oxygen Species (ROS)

Intracellular ROS generation was assessed using 2ʹ,7ʹ-dichlorofluorescein diacetate (DCFH-DA). *Staphylococcus aureus* biofilms were developed in 96-well microtiter plates by incubating overnight cultures for 24 hours, as described above. Following incubation, the preformed biofilms were treated with 100 μM of photosensitizer (TBO or curcumin) in the dark for 30 minutes. Subsequently, 20 µM DCFH-DA was added to each well, and the plates were exposed to white household LED light bulbs for different time intervals. In the light-only control group, biofilms were exposed to LED light in the absence of photosensitizer. After treatment, fluorescence intensity was measured using a Hitachi F-4500X fluorescence spectrophotometer (Tokyo, Japan) equipped with a data-processing unit. The excitation wavelength was set at 485 nm with a slit width of 1.5 nm, and emission from the oxidized DCF was recorded as a measure of ROS levels, as previously described (Misba et al., 2016).

### Live/Dead Staining and Confocal Laser Scanning Microscopy (CLSM)

Confocal laser scanning microscopy (CLSM) was employed to assess the effects of TBO- and curcumin-mediated antimicrobial photodynamic therapy (aPDT) on preformed *Staphylococcus aureus* biofilms following exposure to varying doses of LED light, as described by Shailaja et al. (2022). Biofilms were established by incubating bacterial cultures in glass-bottom confocal dishes (Genetix Biotech Asia) at 37 °C for 24 hours. After incubation, non-adherent cells were carefully removed by washing twice with sterile phosphate-buffered saline (PBS, pH 7.4). Subsequently, the biofilms were treated with 100 µM of either TBO or curcumin and irradiated with white LED light for 5, 10, or 15 minutes. Control groups included untreated biofilms and those exposed to LED light alone (without photosensitizer). Following treatment, the biofilms were stained with a mixture of SYTO 9 and propidium iodide (PI), then incubated at 37 °C for 1 hour in the dark. Fluorescence was visualized using a FluoView FV1000 confocal laser scanning microscope (Olympus, Tokyo, Japan) equipped with argon and HeNe lasers. Excitation wavelengths were set at 488 nm for SYTO 9 and 536 nm for PI to distinguish live and dead cells within the biofilm structure.

### In Vivo Study

#### Chronic Wound Rat Model

A total of 30 adult male Wistar rats, each weighing between 250–300 g, were used to establish the chronic wound model. The animals were randomly divided into six groups, with five rats in each group. Group 1 served as the normal control and consisted of healthy rats without any skin abrasion. Group 2 included rats with skin abrasions that received no treatment (untreated control). Group 3 comprised rats with skin abrasions treated with toluidine blue O (TBO) alone, while Group 4 received TBO treatment followed by exposure to LED light for 15 minutes. Group 5 included rats with skin abrasions treated with curcumin only, and Group 6 received curcumin treatment followed by 15 minutes of LED light exposure.

All treatments were applied topically to the wound area. Throughout the study, animals were housed in cages under standard laboratory conditions with a 12-hour light/dark cycle and provided with food and water ad libitum.

#### Induction of Infection

Rats were anesthetized via intraperitoneal injection of a ketamine/xylazine cocktail. The dorsal surface of each rat was shaved, and chronic wounds were created by drawing 6×6 crossed scratch lines within a 1×1 cm² area using a 28-gauge needle. Five minutes after wounding, each defined area was inoculated with 100 µL of a *Staphylococcus aureus* suspension containing 10⁸ CFU in phosphate-buffered saline (PBS), using a pipette tip. The inoculated wounds were left undisturbed for 24 hours to allow biofilm formation.

#### Treatment via Antimicrobial Photodynamic Therapy

Treatment started on day 2 post-infection. All procedures involving TBO and curcumin were carried out under low-light conditions or in complete darkness, except during the illumination phase. A volume of 100 µL of either TBO or curcumin (100 µM) was applied to the center of the infected wound area and evenly spread across the entire surface. Immediately following application, the treated wounds were exposed to LED light for 15 minutes to initiate photodynamic activation.

#### Bacterial Load Reduction (CFU/mL)

Twenty-four hours post-infection, swab samples were collected from the infected skin surfaces of all experimental groups, including control, untreated, TBO only, TBO with light, curcumin only, and curcumin with light treatment. The collected samples were serially diluted in phosphate-buffered saline and plated on Brain Heart Infusion (BHI) agar to determine the total culturable bacterial count. The plates were incubated at 37 °C for 24 hours, after which *Staphylococcus aureus* colonies were enumerated to assess bacterial load reduction, as described by Gupta et al. (2021).

#### Histopathological Analysis

Fresh skin biopsies were collected from control, untreated, and aPDT-treated rat groups and immediately fixed in 10% phosphate-buffered formalin (pH 7.4). The tissues were dehydrated through a graded series of ethanol, cleared in xylene, and embedded in molten paraplast at 58– 62 °C. To assess the histological response to treatment and microbial presence, serial sections of 5 μm thickness were prepared and stained with hematoxylin and eosin (H&E), following the protocol described by El-Gayar et al. (2022).

#### Cytokine Assays

Blood samples were collected from anesthetized rats via retro-orbital bleeding using sterile collection tubes. Samples were obtained from all experimental groups, including control, untreated, TBO-only treated, TBO with light, curcumin-only treated, and curcumin with light groups to assess cytokine levels. Immediately after blood collection, all animals were euthanized in accordance with the approved protocol. The collected blood samples were centrifuged at 5000 rpm for 5 minutes, and the resulting serum supernatant was aliquoted into sterile microcentrifuge tubes and stored at −80 °C for further analysis, as described by Karamese et al. (2016). Cytokine levels were quantified using enzyme-linked immunosorbent assay (ELISA) kits according to the manufacturers’ instructions. The following kits were used: Rat IL-6 ELISA Kit (RayBiotech, Cat. No. ELR-IL6), Rat IL-1β ELISA Kit (Elabscience, Cat. No. E-EL-R0012), and Rat IFN-γ ELISA Kit (Elabscience, Cat. No. E-EL-R0009). Absorbance readings were obtained using a Bio-Rad Microplate Reader (India), and levels of IL-6, IL-1β, and IFN-γ were calculated accordingly.

### Statistical Analysis

Statistical comparisons among multiple groups were performed using one-way analysis of variance (ANOVA). Data are presented as mean ± standard deviation (SD) from three independent experiments. A p-value of less than 0.05 was considered statistically significant, as described by Kim (2017).

## RESULTS

### Antibacterial Activity of TBO and Curcumin-Mediated Photodynamic Therapy (PDT) Against Planktonic S. aureus

The antibacterial efficacy of TBO and curcumin, both with and without household LED light exposure, was assessed using a colony formation assay. After 24 hours of incubation, a reduction of *S. aureus* by 0.18 log₁₀ CFU/ml was observed in the group exposed to LED light alone. The TBO-only treatment resulted in a 0.43 log₁₀ CFU/ml reduction. When TBO treatment was followed by LED light exposure for 5 and 10 minutes, the bacterial reduction increased to 1.13 log₁₀ CFU/ml and 4.85 log₁₀ CFU/ml, respectively. Notably, no bacterial growth was detected when the plates were exposed to LED light for 15 minutes **(Figure S1. a)**. Similarly, in the curcumin-treated groups, a 0.26 log₁₀ CFU/ml reduction was observed with LED light exposure alone, while curcumin treatment alone resulted in a 0.29 log₁₀ CFU/ml reduction. When curcumin treatment was followed by LED light exposure for 5, 10, and 15 minutes, the bacterial reduction was 0.72 log₁₀ CFU/ml, 0.9 log₁₀ CFU/ml, and 2.81 log₁₀ CFU/ml, respectively **(Figure S1. b)**.

### Reduction of Extracellular Polymeric Substance (EPS) Production in S. aureus Biofilm

EPS, a crucial component for biofilm stability, was quantified using the Congo red binding assay. Treatment with TBO alone resulted in a 5.66% reduction in EPS production. However, when TBO treatment was combined with LED light exposure, the EPS reduction increased to 32.08%, 69.8%, and 96.22% for 5, 10, and 15 minutes of exposure, respectively **(Figure 1a)**. Curcumin treatment alone resulted in a 6.12% reduction in EPS production. When followed by LED light exposure for 5, 10, and 15 minutes, the reduction increased to 16.32%, 24.48%, and 48.98%, respectively **(Figure 1b)**.

**Figure 1.**
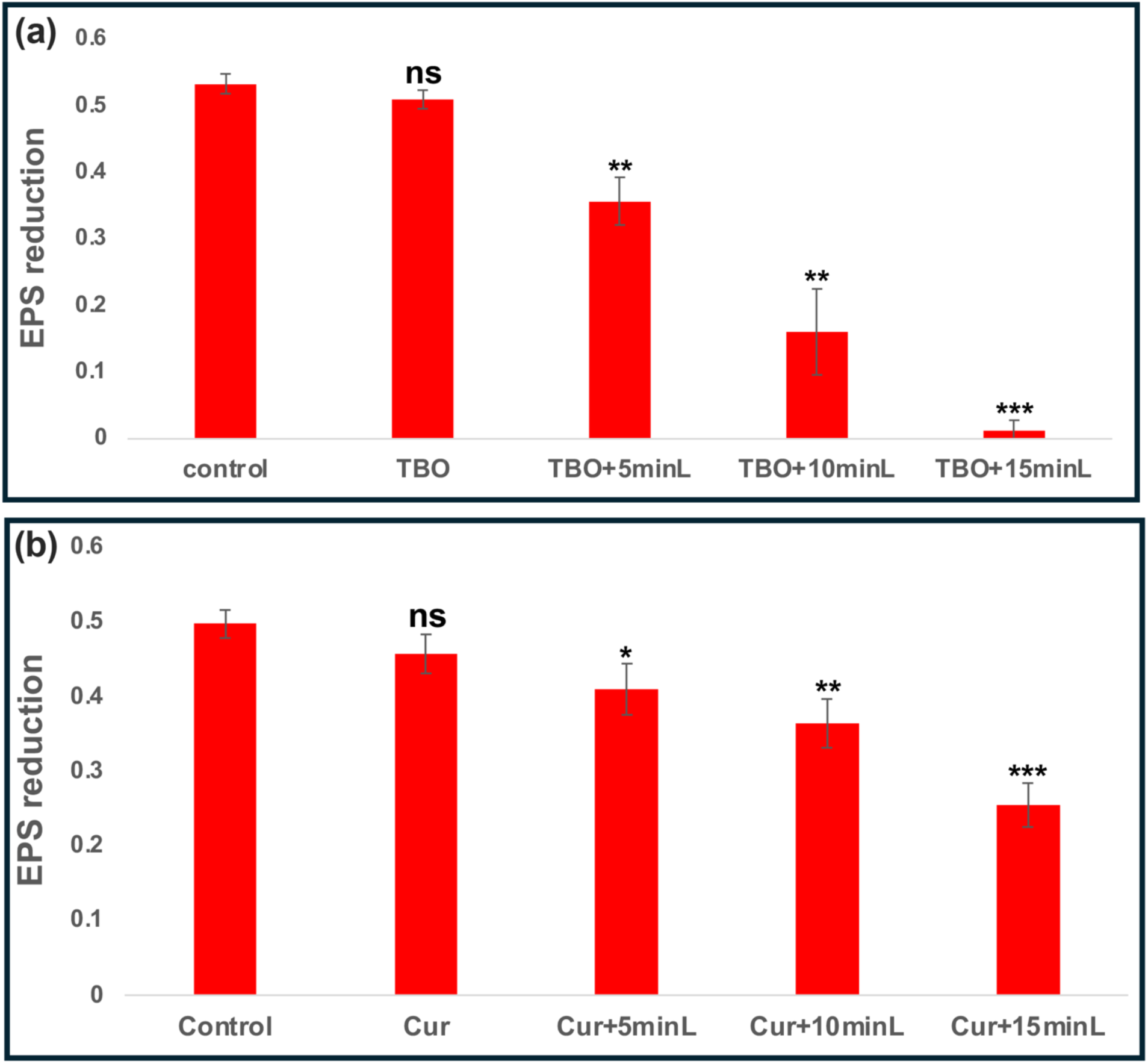
Quantification of extracellular polymeric substances (EPS) in *S. aureus* biofilms following treatment with (a) TBO and (b) curcumin, assessed via the Congo Red assay. Absorbance was recorded at 490 nm. Values represent mean ± SD (n = 3). Comparisons with the control group were performed using Student’s *t*-test, and multiple group comparisons were analyzed by one-way ANOVA. Statistical significance: *p* < 0.05 (*), p < 0.01 (), p < 0.001 (*); ns = not significant.

### Intracellular ROS Generation in S. aureus Biofilm

Reactive oxygen species (ROS) production was significantly enhanced following treatment with TBO or curcumin in combination with LED light exposure, compared to treatment with TBO or curcumin alone. No significant ROS generation was observed in the group exposed to LED light alone. Moreover, ROS levels increased progressively with prolonged LED light exposure, with the highest ROS generation observed at 15 minutes **(Figure 2)**.

**Figure 2.**
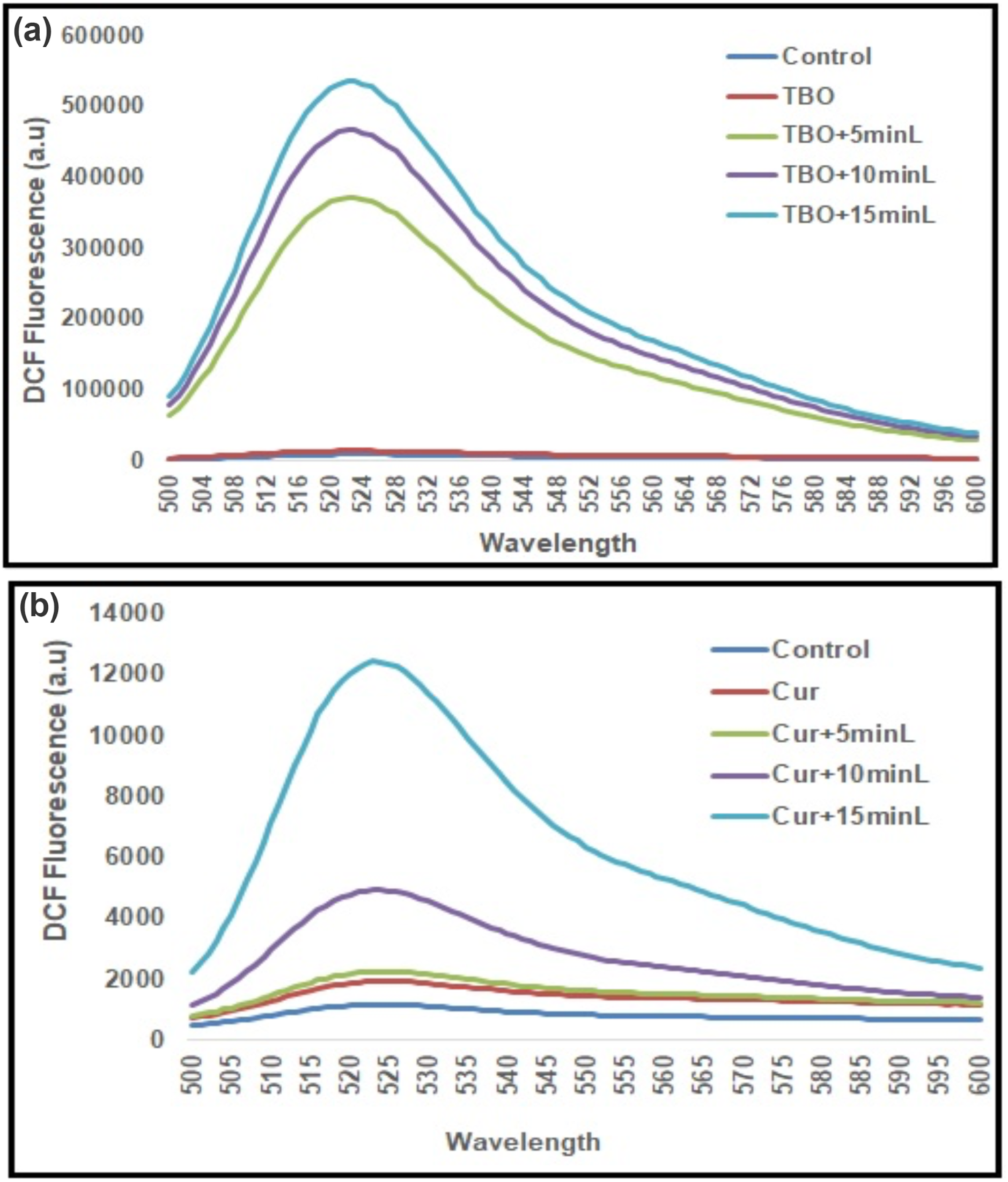
Total reactive oxygen species (ROS) generation following (a) TBO- and (b) curcumin-mediated antimicrobial photodynamic therapy (aPDT) against *S. aureus*, determined by fluorescence intensity of 2′,7′-dichlorofluorescein (DCF). Control represents untreated samples.

### Live/Dead Cell Analysis Using Confocal Laser Scanning Microscopy (CLSM)

CLSM was used to visualize biofilm architecture following TBO- and curcumin-mediated PDT in *S. aureus* biofilms. Biofilms treated with only LED light or left untreated exhibited a uniform green fluorescence, indicating the presence of live cells. In contrast, biofilms treated with TBO or curcumin and subsequently exposed to LED light showed both green and red fluorescence, indicating a mix of live and dead cells. The intensity of red fluorescence, representing dead cells, increased with prolonged LED light exposure, with 15 minutes of exposure yielding the highest proportion of dead cells **(Figure 3)**.

**Figure 3.**
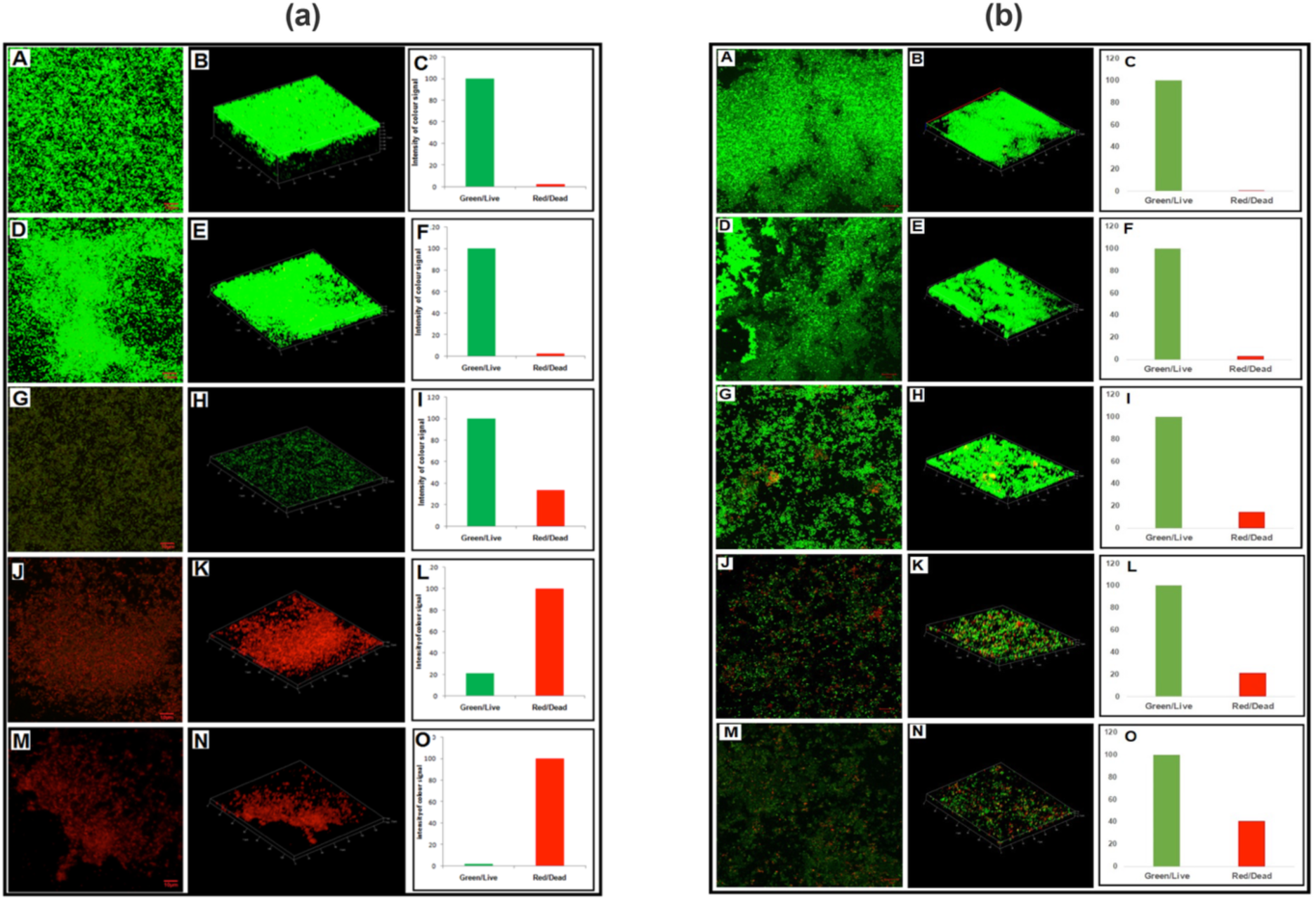
Live/dead cell viability assay of *S. aureus* biofilms post-photodynamic therapy (PDT) with (a) TBO and (b) curcumin. Biofilm cells were stained with SYTO 9 (green, live cells) and propidium iodide (PI; red, dead cells). Scale bar = 10 μm.

### In Vivo Efficacy of TBO- and Curcumin-Mediated PDT in Treating Skin Abrasion

The therapeutic potential of TBO- and curcumin-mediated PDT was evaluated in Wistar rats with skin abrasions **(Figure 4)**. TBO treatment followed by LED light exposure for 15 minutes significantly reduced *S. aureus* colonization within four days. On day four, bacterial load was 2.09 × 10⁴ CFU/ml in the untreated group, 1.5 × 10⁴ CFU/ml in the TBO-treated group, and 1.7 × 10³ CFU/ml in the TBO-treated group exposed to LED light. Similarly, in curcumin- treated groups, bacterial load was 1.3 × 10⁴ CFU/ml, whereas curcumin treatment followed by LED light exposure reduced the bacterial load to 6.2 × 10³ CFU/ml **(Figure S2)**.

**Figure 4.**
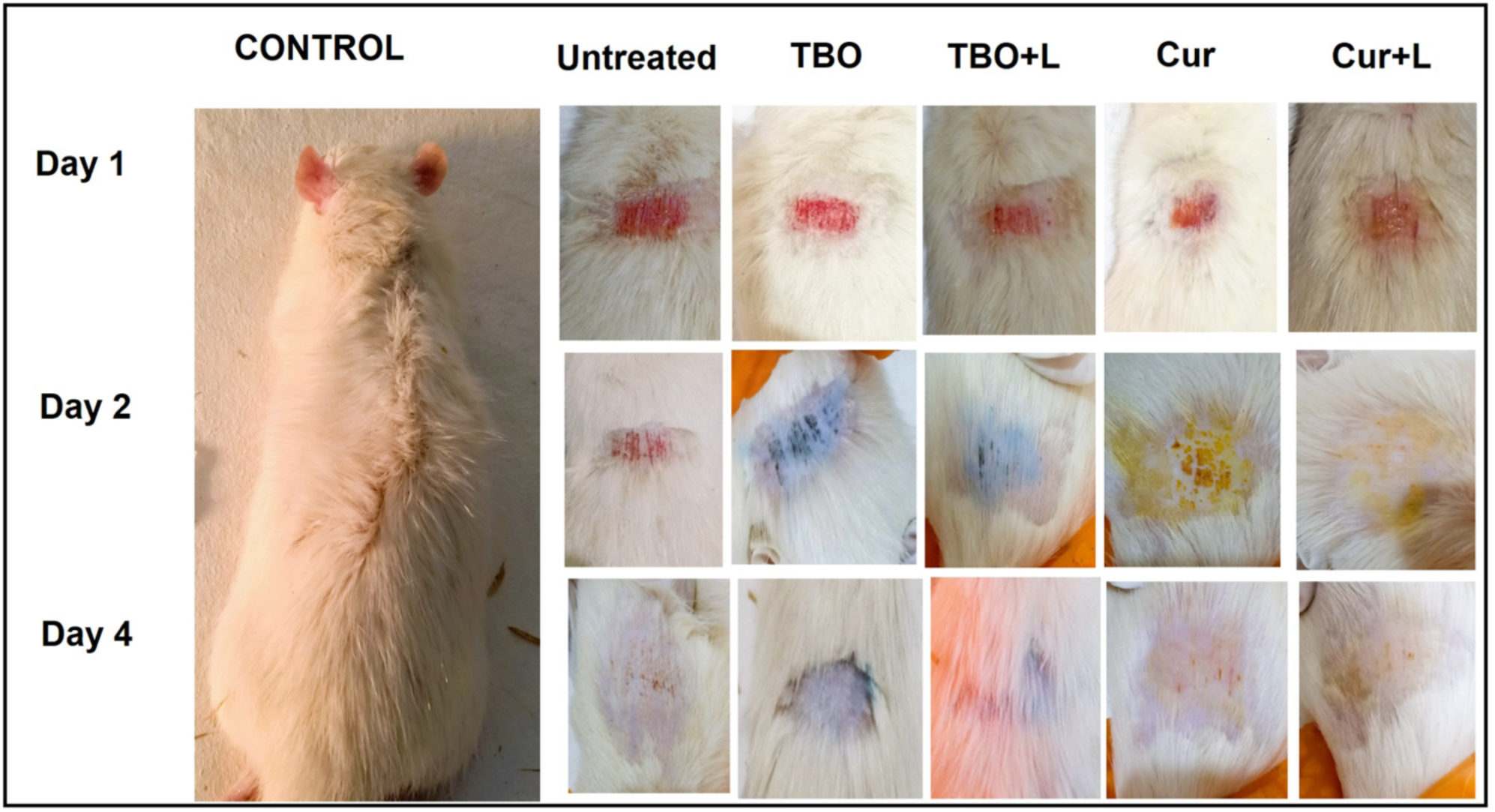
Male Wistar rat skin abrasion model treated with TBO, or curcumin followed by 15 min of LED light exposure.

### Histological Examination of Skin Tissue

Histological examination of the control group revealed normal skin architecture with intact sebaceous glands. In contrast, skin tissues from the infected, untreated rats exhibited dense neutrophilic infiltration and dispersed inflammatory cells, confirming the presence of infection. Rats treated with TBO or curcumin alone showed moderate neutrophilic infiltration. However, rats treated with TBO or curcumin followed by LED light exposure displayed stratified squamous epithelium with sparsely dispersed neutrophils and capillaries, suggesting improved wound healing and reduced inflammation **(Figure 5a)**.

**Figure 5.**
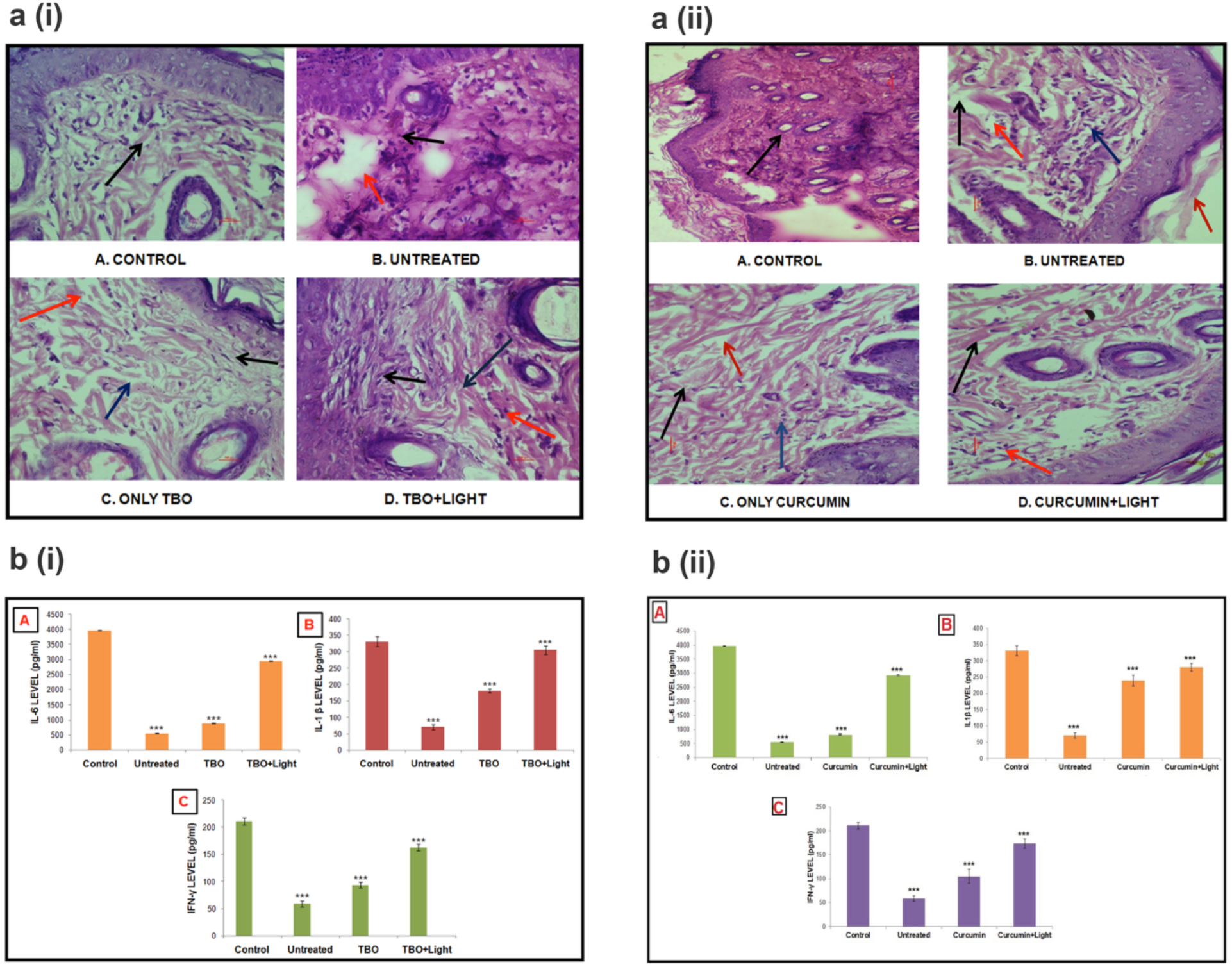
(a) Histopathological evaluation of rat skin tissue using hematoxylin and eosin (H/E) staining after treatment with (i) TBO or (ii) curcumin, assessing tissue response to microbial infection. Scale bar = 500 μm. (b) Anti-inflammatory effects of (i) TBO- and (ii) curcumin- mediated aPDT in the rat skin abrasion model, determined by quantifying cytokine levels: (A) IL-6, (B) IL-1β, and (C) IFN-γ.

### Effect of TBO- and Curcumin-Mediated PDT on Inflammatory Cytokines

The expression of pro-inflammatory cytokines (IL-6, IL-1β, and IFN-ɣ) was analysed across different experimental groups. Compared to untreated rats and those treated with PS (TBO or curcumin) alone, rats treated with PS followed by LED light exposure for 15 minutes exhibited significantly elevated levels of IL-6, IL-1β, and IFN-ɣ, indicating an immune response triggered by PDT **(Figure 5b)**.

## DISCUSSION

Antimicrobial resistance (AMR) has emerged as a major global health concern, necessitating the urgent development of alternative therapeutic strategies (Martinez et al. 2014). Multidrug-resistant bacterial strains, including *Pseudomonas aeruginosa, Staphylococcus aureus, Acinetobacter baumannii, Escherichia coli,* and others, are responsible for severe infections ranging from bacteremia and meningitis to skin, pulmonary, and urinary tract infections, as well as prosthetic device-related complications (Taylor et al. 2023). The increasing resistance of these pathogens to conventional antibiotics underscores the need for alternative approaches (Munita et al. 2016; Afzal et al. 2022). One promising method is photodynamic therapy (PDT), which employs photosensitizers (PS) activated by light to generate reactive oxygen species (ROS), leading to bacterial cell death (Hu et al. 2023; Akhtar et al. 2024; Khan et al. 2020).

The antimicrobial photoinactivation of *S. aureus* using toluidine blue O (TBO) and curcumin has been previously explored (He et al. 2022; Jiang et al. 2014; Akhtar et al. 2021; Akhtar et al.2021). In this study, we aimed to evaluate the antibacterial efficacy of TBO and curcumin upon irradiation with household LED light as a non-toxic, anti-proliferative alternative for combating multidrug-resistant *S. aureus*.

The colony-forming unit (CFU) assay demonstrated enhanced antibacterial activity of TBO and curcumin against *S. aureus* in the presence of LED light, with a more pronounced effect observed with increased exposure time. Notably, complete bacterial eradication was achieved in plates treated with TBO followed by 15 minutes of LED light exposure, indicating its superior photosensitizing properties compared to curcumin **(Figure S1)**.

Exopolysaccharide (EPS) production, a key factor in biofilm stability, was assessed using the Congo red assay (Kaplan et al. 2018). A significant reduction in EPS production was observed in groups treated with TBO or curcumin in combination with LED light, as compared to treatment with photosensitizers alone or the control. The extent of EPS reduction correlated positively with increasing LED light exposure time, with TBO exhibiting greater efficacy than curcumin even at shorter exposure durations. This further reinforces the superior photosensitizing ability of TBO **(Figure 1)**.

To elucidate the underlying antibacterial mechanism, we quantified ROS generation using the DCFH-DA assay. The results revealed significantly higher ROS levels in groups treated with photosensitizers followed by LED light exposure compared to those treated with photosensitizers alone, light alone, or the control **(Figure 2)**. Moreover, ROS production increased with longer light exposure durations, with TBO again demonstrating greater efficacy than curcumin. These findings align with the CFU and Congo red assay results, confirming the enhanced antibacterial activity of PDT.

Morphological alterations in bacterial biofilms were further analyzed using confocal laser scanning microscopy (CLSM) (Vassena et al. 2014). The disruption of *S. aureus* biofilm was evident in groups treated with TBO or curcumin followed by LED light irradiation, whereas the control and light-only groups exhibited intact biofilms **(Figure 3)**. Biofilm disruption and bacterial cell death increased proportionally with extended LED light exposure. The ratio of dead to live cells was markedly higher in TBO-treated groups compared to curcumin-treated groups, further validating its superior photosensitizing activity.

The *in vivo* efficacy of TBO and curcumin-mediated PDT was assessed in a rat skin abrasion model **(Figure 4)**. The results demonstrated a significant reduction in *S. aureus* colonization in groups treated with TBO or curcumin followed by LED light exposure, compared to untreated groups or those treated with photosensitizers alone **(Figure S2)**. Histopathological analysis further confirmed improved tissue healing in groups receiving PDT, as indicated by reduced neutrophilic infiltration and enhanced epithelial regeneration, compared to untreated groups or those receiving photosensitizers alone **(Figure 5a)**.

To investigate the role of inflammatory response in wound healing, we analysed the expression levels of pro-inflammatory cytokines IL-6, IL-1β, and IFN-ɣ. A significant upregulation of these cytokines was observed in groups treated with PDT compared to untreated or photosensitizer-only groups **(Figure 5b)**. During the early stages of infection, inflammation plays a crucial role in bacterial clearance and wound healing. The observed increase in cytokine expression in PDT-treated groups suggests an early inflammatory response, which is typically followed by a resolution phase, facilitating tissue repair and restoration (Guan et al. 2022).

Overall, our findings demonstrate the potent antibacterial and wound-healing efficacy of TBO and curcumin-mediated PDT using household LED light. The data highlight the potential of PDT as a promising alternative therapeutic approach for combating multidrug-resistant *S. aureus* infections, with TBO exhibiting superior efficacy over curcumin. Future studies should focus on optimizing treatment parameters and exploring PDT applications in clinical settings for enhanced antimicrobial and wound-healing outcomes.

## CONCLUSIONS

This study establishes antimicrobial photodynamic therapy (PDT) using toluidine blue O (TBO) and curcumin, activated by a readily available household LED light, as a powerful and accessible approach for combating multidrug-resistant *Staphylococcus aureus* infections. Both photosensitizers demonstrated potent antibacterial activity, with TBO exhibiting superior efficacy in bacterial eradication, exopolysaccharide reduction, and biofilm disruption. The marked increase in reactive oxygen species (ROS) generation under LED illumination underscores the mechanistic basis of the enhanced antibacterial effects. *In vivo* experiments further confirmed accelerated wound healing and substantial reduction in bacterial load in a rat skin abrasion model, emphasizing the clinical relevance of this strategy. These findings position household LED–mediated PDT, particularly with TBO, as a cost-effective, non-toxic, and scalable alternative to conventional antibiotics, with strong potential for translation into real-world therapeutic applications against multidrug-resistant pathogens.

## Author contributions

Conceptualization, L.M., F.A., and A.U.K.; methodology, L.M., F.A., and A.U.K.; formal analysis, L.M., F.A., and A.U.K.; investigation, L.M. and F.A.; resources, A.U.K.; writing-original L.M., F.A., and S.M.; writing-review and editing, L.M., F.A., S.M., and A.U.K.; visualization, L.M., F.A., S.M., and A.U.K.; supervision, A.U.K.; and funding acquisition, A.U.K.

## Notes

The authors declare no competing financial interest.

## ACKNOWLEDGMENTS

The authors would also like to acknowledge university sophisticated instruments facility (USIF), AMU, for providing instrumental support.

## REFERENCES

Abrahamse, H., & Hamblin, M. R. (2016). New photosensitizers for photodynamic therapy. Biochemical Journal, 473(4), 347–364. 10.1042/BJ20150942

Aebisher, D., Przygórzewska, A., & Bartusik-Aebisher, D. (2024). Natural photosensitizers in clinical trials. Applied Sciences, 14, 8436. 10.3390/app14188436

Afzal, O., Altamimi, A. S. A., Nadeem, M. S., Alzarea, S. I., Almalki, W. H., Tariq, A., Mubeen, B., Murtaza, B. N., Iftikhar, S., Kazmi, I. (2022). Nanoparticles in drug delivery: From history to therapeutic applications. Nanomaterials, 12, 4494. 10.3390/nano12244494

Akhtar, F., Khan, A. U., Qazi, B., Kulanthaivel, S., Mishra, P., Akhtar, K., & Ali, A. (2021). A nano phototheranostic approach of toluidine blue conjugated gold-silver core shells mediated photodynamic therapy to treat diabetic foot ulcer. Scientific Reports, 11(1), 24464. 10.1038/s41598-021-04008-x

Akhtar, F., Khan, A. U., Misba, L., & Akhtar, K. (2024). The dual role of photodynamic therapy to treat cancer and microbial infection. Drug Discovery Today, 104099.

Akhtar, F., Khan, A. U., Misba, L., Akhtar, K., & Ali, A. (2021). Antimicrobial and antibiofilm photodynamic therapy against vancomycin-resistant *Staphylococcus aureus* (VRSA) induced infection in vitro and in vivo. European Journal of Pharmaceutics and Biopharmaceutics, 160, 65–76.

Akhtar, F., & Khan, A. U. (2021). Antimicrobial photodynamic therapy (aPDT) against vancomycin-resistant *Staphylococcus aureus* (VRSA) biofilm disruption: A putative role of phagocytosis in infection control. Photodiagnosis and Photodynamic Therapy, 36, 102552.

ANOVA: Analysis of Variance between groups. Retrieved from www.physics.csbsju.edu/stats/anova.html

Centers for Disease Control and Prevention. (2019, November 6). The biggest antibiotic-resistant threats in the U.S. Retrieved from https://www.cdc.gov/drugresistance/threat-report-2019.html

Centers for Disease Control and Prevention. (2022). COVID-19: U.S. Impact on antimicrobial resistance, special report 2022. U.S. Department of Health and Human Services, CDC. Retrieved from https://www.cdc.gov/drugresistance/covid19.html

Correia, J. H., Rodrigues, J. A., Pimenta, S., Dong, T., & Yang, Z. (2021). Photodynamic therapy review: Principles, photosensitizers, applications, and future directions. Pharmaceutics, 13(9), 1332. 10.3390/pharmaceutics13091332

Denissen, J., Reyneke, B., Waso-Reyneke, M., et al. (2022). Prevalence of ESKAPE pathogens in the environment: Antibiotic resistance status, community-acquired infection, and risk to human health. International Journal of Hygiene and Environmental Health, 244, 114006.

Dincer, S., Uslu, F. M., & Delik, A. (2020). Antibiotic resistance in biofilm. In Bacterial Biofilms. IntechOpen. 10.5772/intechopen.92388

El-Gayar, M. H., Ishak, R. A. H., Esmat, A., Aboulwafa, M. M., & Aboshanab, K. M. (2022). Evaluation of lyophilized royal jelly and garlic extract emulgels using a murine model infected with methicillin-resistant *Staphylococcus aureus*. AMB Express, 12(1),37. 10.1186/s13568-022-01378-x

Escudero, A., Carrillo-Carrión, C., Castillejos, M. C., Romero-Ben, E., Rosales-Barrios, C., & Khiar, N. (2021). Photodynamic therapy: Photosensitizers and nanostructures. Materials Chemistry Frontiers, 5, 3788–3812. 10.1039/D0QM00922A

Ghorbani, J., Rahban, D., Aghamiri, S., Teymouri, A., & Bahador, A. (2018). Photosensitizers in antibacterial photodynamic therapy: An overview. Laser Therapy, 27(4), 293–302. 10.5978/islsm.27_18-RA-01

Guan, Y., Liu, N., Yu, Y., Zhou, Q., Chang, M., Wang, Y., & Yao, S. (2022). Pathological comparison of rat pulmonary models induced by silica nanoparticles and indium-tin oxide nanoparticles. International Journal of Nanomedicine, 17, 4277– 4292. 10.2147/IJN.S380259

Gupta, V., & Tyagi, A. (2021). A rat model of polymicrobial infection in full-thickness excision wounds. Journal of Tissue Viability, 30(4), 537–543. 10.1016/j.jtv.2021.06.003

He, Y., Pang, J., Yang, Z., Zheng, M., Yu, Y., Liu, Z., Zhao, B., Hu, G., & Yin, R. (2022). Toluidine blue O-induced photoinactivation inhibits the biofilm formation of methicillin-resistant *Staphylococcus aureus*. Photodiagnosis and Photodynamic Therapy, 39, 102902. 10.1016/j.pdpdt.2022.102902

Hu, X., et al. (2023). Antimicrobial photodynamic therapy encapsulation technology: Frontier exploration and application prospects of novel antimicrobial technology. Chemical Engineering Journal, 477, 146773.

Ikuta, K. S., Swetschinski, L. R., Robles Aguilar, G., et al. (2022). Global mortality associated with 33 bacterial pathogens in 2019: A systematic analysis for the Global Burden of Disease Study 2019. The Lancet, 400, 2221–2248.

Jiang, Y., Leung, A. W., Hua, H., Rao, X., & Xu, C. (2014). Photodynamic action of LED-activated curcumin against *Staphylococcus aureus* involving intracellular ROS increase and membrane damage. International Journal of Photoenergy, 2014, 637601. 10.1155/2014/637601

Kaplan, J. B., Mlynek, K. D., Hettiarachchi, H., Alamneh, Y. A., Biggemann, L., Zurawski, D. V., Black, C. C., Bane, C. E., Kim, R. K., & Granick, M. S. (2018). Extracellular polymeric substance (EPS)-degrading enzymes reduce staphylococcal surface attachment and biocide resistance on pig skin in vivo. PLOS One, 13(10), e0205526. 10.1371/journal.pone.0205526

Karamese, M., Erol, H. S., Albayrak, M., Findik Guvendi, G., Aydin, E., & Aksak Karamese, S. (2016). Anti-oxidant and anti-inflammatory effects of apigenin in a rat model of sepsis: An immunological, biochemical, and histopathological study. Immunopharmacology and Immunotoxicology, 38(3), 228–237.

Khan, S., Khan, S. N., Akhtar, F., Misba, L., Meena, R., & Khan, A. U. (2020). Inhibition of multi-drug resistant *Klebsiella pneumoniae*: Nanoparticles induced photoinactivation in presence of efflux pump inhibitor. European Journal of Pharmaceutics and Biopharmaceutics, 157, 165–174.

Liu, H. Y., Prentice, E. L., & Webber, M. A. (2024). Mechanisms of antimicrobial resistance in biofilms. npj Antimicrobial Resistance, 2, 27. 10.1038/s44259-024-00046-3

Martinez, J. L. (2014). General principles of antibiotic resistance in bacteria. Drug Discovery Today: Technologies, 11, 33–39. 10.1016/j.ddtec.2014.02.001

Mathur, A., Parihar, A. S., Modi, S., & Kalra, A. (2023). Photodynamic therapy for ESKAPE pathogens: An emerging approach to combat antimicrobial resistance (AMR). Microbial Pathogenesis. 10.1016/j.micpath.2023.106307

Mazzariol, A., Bazaj, A., & Cornaglia, G. (2017). Multi-drug-resistant Gram-negative bacteria causing urinary tract infections: A review. Journal of Chemotherapy, 29, 2–9.

Minasyan, H. (2017). Sepsis and septic shock: Pathogenesis and treatment perspectives. Journal of Critical Care, 40, 229–242.

Misba, L., & Khan, A. U. (2018). Enhanced photodynamic therapy using light fractionation against *Streptococcus mutans* biofilm: Type I and type II mechanism. Future Microbiology, 13, 437–454. 10.2217/fmb-2017-0207

Misba, L., Kulshrestha, S., & Khan, A. U. (2016). Antibiofilm action of a toluidine blue O-silver nanoparticle conjugate on *Streptococcus mutans*: A mechanism of type I photodynamic therapy. Biofouling, 32(3), 313–328. 10.1080/08927014.2016.1141899

Munita, J. M., & Arias, C. A. (2016). Mechanisms of antibiotic resistance. Microbiology Spectrum, 4. 10.1128/microbiolspec.VMBF-0016-2015

Murray, C. J. L., Ikuta, K. S., Sharara, F., et al. (2022). Global burden of bacterial antimicrobial resistance in 2019: A systematic analysis. The Lancet, 399, 629–655.

Pemula, G., Girigoswami, K., & Muthukrishnan, S. (2022). Nanoformulation of tetrapyrrole derivatives in photodynamic therapy: A focus on bacteriochlorin. Evidence-Based Complementary and Alternative Medicine, 3011918, 12 pages. 10.1155/2022/3011918

Shailaja, A., Bruce, T. F., Gerard, P., Powell, R. R., Pettigrew, C. A., & Kerrigan, J. L. (2022). Comparison of cell viability assessment and visualization of *Aspergillus niger* biofilm with two fluorescent probe staining methods. Biofilm, 4, 100090. 10.1016/j.bioflm.2022.100090

Sharma, D., Misba, L., & Khan, A. U. (2019). Antibiotics versus biofilm: An emerging battleground in microbial communities. Antimicrobial Resistance and Infection Control, 8, 76. 10.1186/s13756-019-0533-3

Singh, R., Dwivedi, S. P., Gaharwar, U. S., et al. (2020). Recent updates on drug resistance in *Mycobacterium tuberculosis*. Journal of Applied Microbiology, 128, 1547–1567.

Taylor, T. A., & Unakal, C. G. (2023, July 17). Staphylococcus aureus infection. In StatPearls [Internet]. Treasure Island (FL): StatPearls Publishing. Available from: https://www.ncbi.nlm.nih.gov/books/NBK441868/

Terreni, M., Taccani, M., & Pregnolato, M. (2021). New antibiotics for multidrug-resistant bacterial strains: Latest research developments and future perspectives. Molecules, 26, 2671.

Vassena, C., Fenu, S., Giuliani, F., Fantetti, L., Roncucci, G., Simonutti, G., Romanò, C. L., De Francesco, R., & Drago, L. (2014). Photodynamic antibacterial and antibiofilm activity of RLP068/Cl against *Staphylococcus aureus* and *Pseudomonas aeruginosa* forming biofilms on prosthetic material. International Journal of Antimicrobial Agents, 44(1), 47–55. 10.1016/j.ijantimicag.2014.03.012

Venkateswaran, P., Vasudevan, S., David, H., et al. (2023). Revisiting ESKAPE pathogens: Virulence, resistance, and combating strategies focusing on quorum sensing. Frontiers in Cellular and Infection Microbiology, 13.

