## Supplementary material for "Household LED Light-Mediated Photodynamic Therapy for Effective Inhibition of Multidrug-Resistant *Staphylococcus aureus*": Figure S1.a, Figure S1.b, Figure S2,

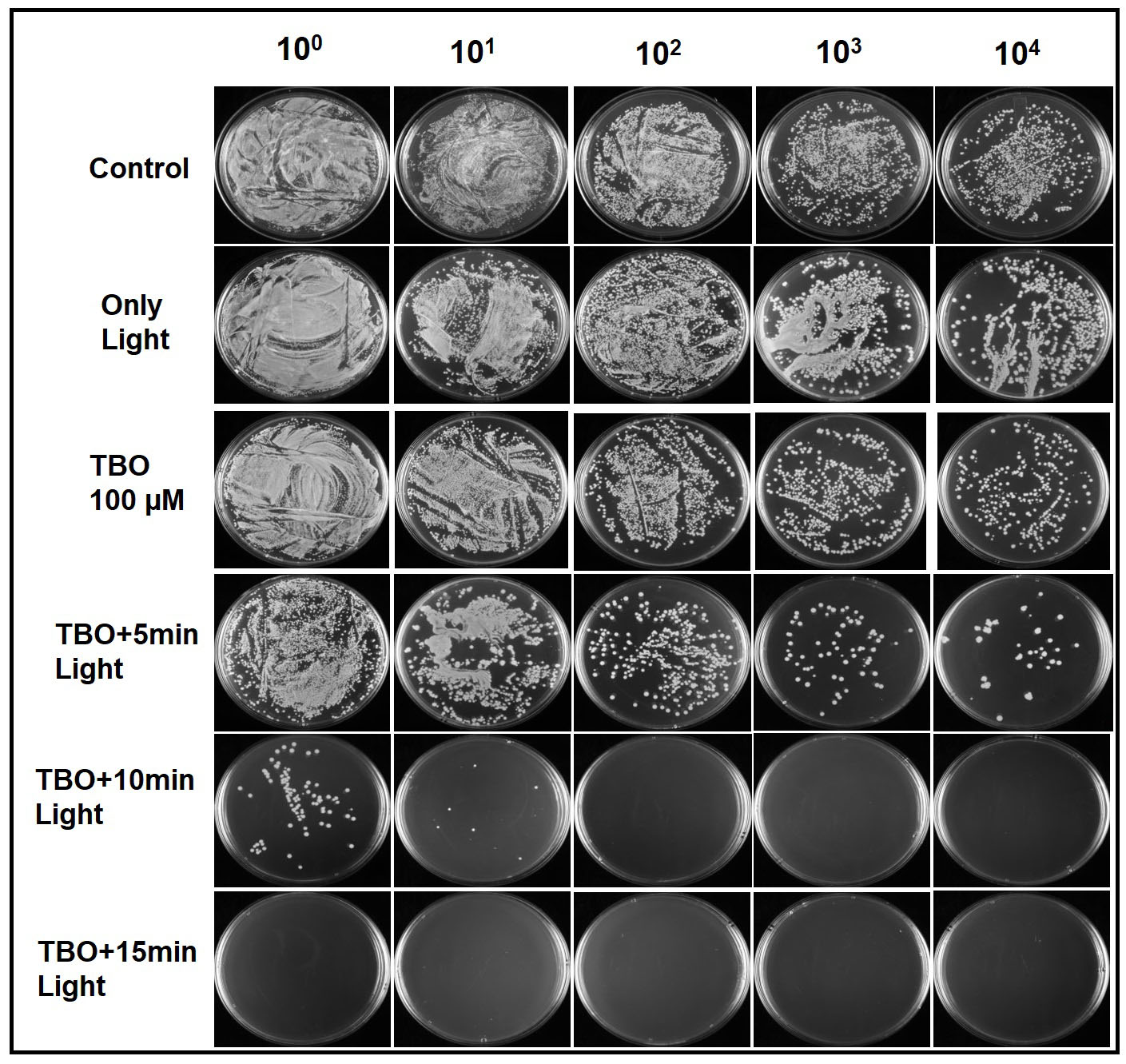

**Figure S1. a** Antibacterial activity of TBO-mediated photodynamic therapy (PDT) against S. aureus assessed by colony-forming unit (CFU) assay. Bacterial cultures were treated with TBO (100 μM) followed by LED light exposure for 5-, 10-, or 15-min. Serial dilutions (10⁰-10⁴) were plated on agar, and representative plates are shown for each treatment condition, including controls (untreated, light only, and TBO without light).

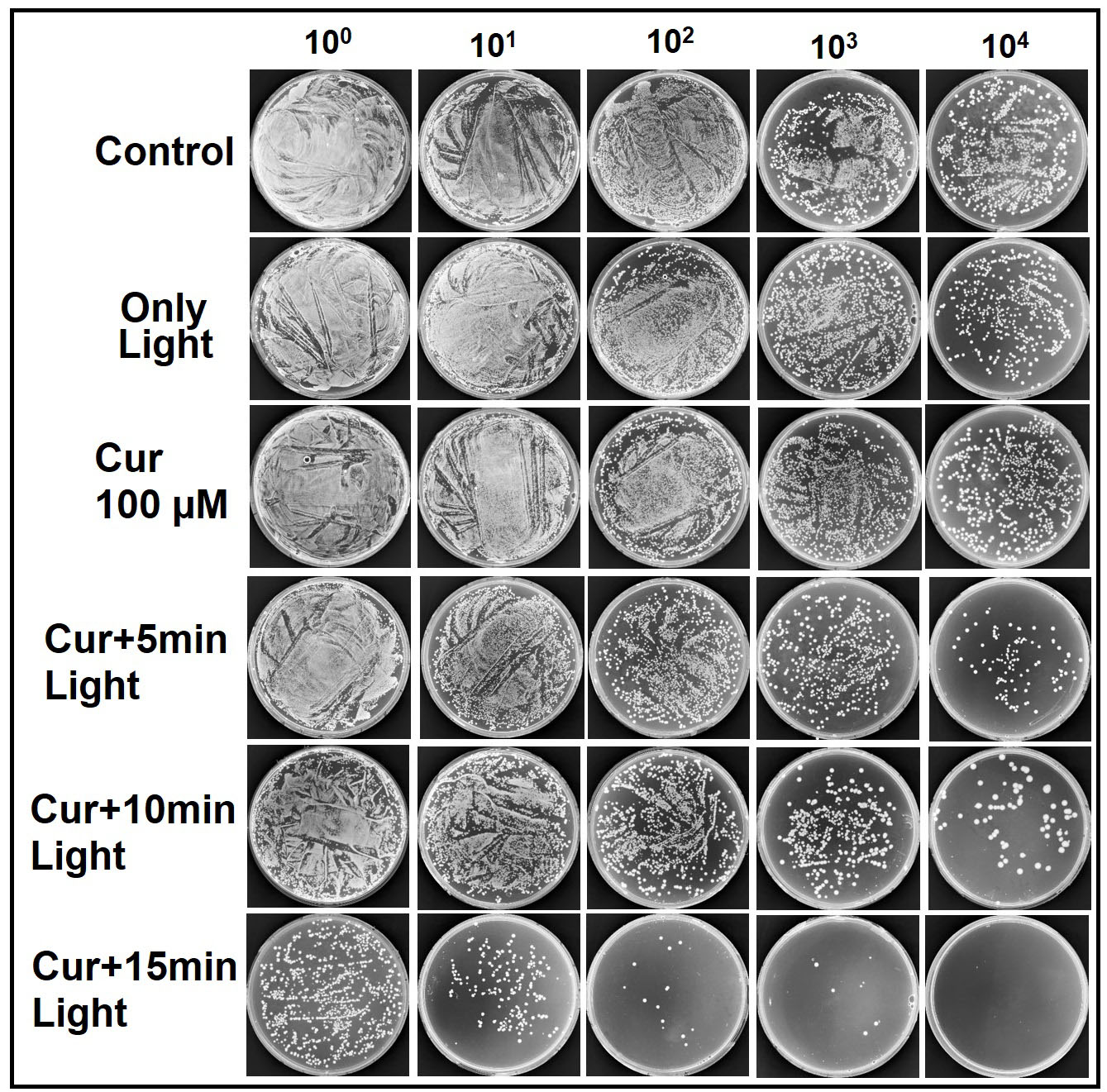

**Figure S1. b** Antibacterial activity of curcumin-mediated photodynamic therapy (PDT) against S. aureus assessed by colony-forming unit (CFU) assay. Bacterial cultures were treated with curcumin (100 μM) followed by LED light exposure for 5-, 10-, or 15-min. Serial dilutions (10⁰-10⁴) were plated on agar, and representative plates are shown for each treatment condition, including controls (untreated, light only, and curcumin without light).

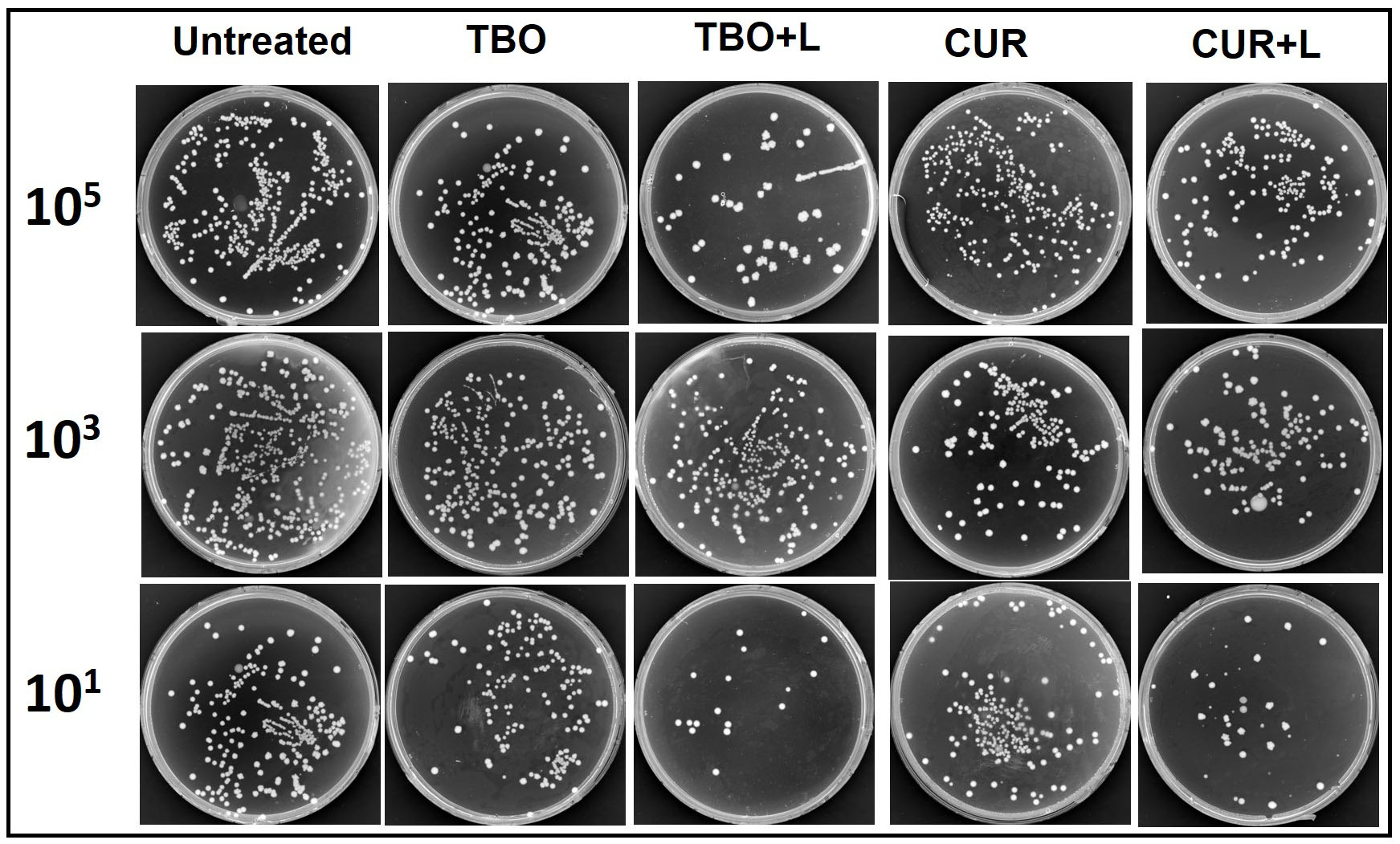

**Figure S2.**  Effect of TBO- and curcumin-mediated photodynamic therapy (PDT) on S. aureus colonization *in vivo*. Colony-forming unit (CFU) assay of bacterial load from skin samples collected four days post-treatment. Groups include untreated control, TBO alone, TBO + LED light (15 min), curcumin alone, and curcumin + LED light (15 min). Serial dilutions (10⁵, 10³, and 10¹) were plated on agar, and representative plates are shown.
